# Identification of Kinome Biomarkers and Therapeutic Rescue in cGMP-Linked Retinal Degeneration

**DOI:** 10.64898/2026.11.06.763903

**Authors:** Iskalen Cansu Topcu Okan, Mehri Ahmadian, Halit Yusuf Altay, Simar Pal Singh, Ghazaleh Kheirandish, Samed Ozer, Pieter Gaillard, Richard Bouffard, Albert Neutzner, Kerem Esmen, Markus Tschopp, Matthew Lawrence, Muhammed Kasim Diril, François Paquet-Durand, Cavit Agca

## Abstract

Inherited retinal degenerations, such as *Retinitis Pigmentosa* (RP) and Leber’s congenital amaurosis (LCA), are major causes of progressive and irreversible vision loss. The large number of genes associated with these disorders makes their clinical and molecular characterization difficult, while systemic biomarkers for disease progression or treatment response remain limited. These challenges were addressed by targeting cGMP-associated mechanisms shared by RP and LCA and integrating systemic biomarker profiling with retinal outcomes. Kinase activity analysis (kinome analysis) in peripheral blood mononuclear cells distinguished healthy and diseased states and captured therapy-associated signatures in *rd10* mice treated with LP-CN03, a liposomal formulation of the cGMP analog CN03. This first systemic analysis in a therapeutically responsive animal model highlights kinome analysis as a potential translational readout. Given shared cGMP signaling in RP and *AIPL1*-associated LCA, LP-CN03’s therapeutic potential was further evaluated in a new CRISPR-Cas9-generated *Aipl1*^⁻/⁻^ mouse model. Treatment with LP-CN03 preserved retinal structure and function, as shown by electroretinography, longitudinal optical coherence tomography, fundus imaging, optomotor testing, and photoreceptor-specific gene expression. Together, these findings provide independent *in vivo* evidence for the neuroprotective effect of LP-CN03 in both cGMP-related RP and LCA, also highlighting the translational value of kinome profiling.

**Graphical Abstract:** 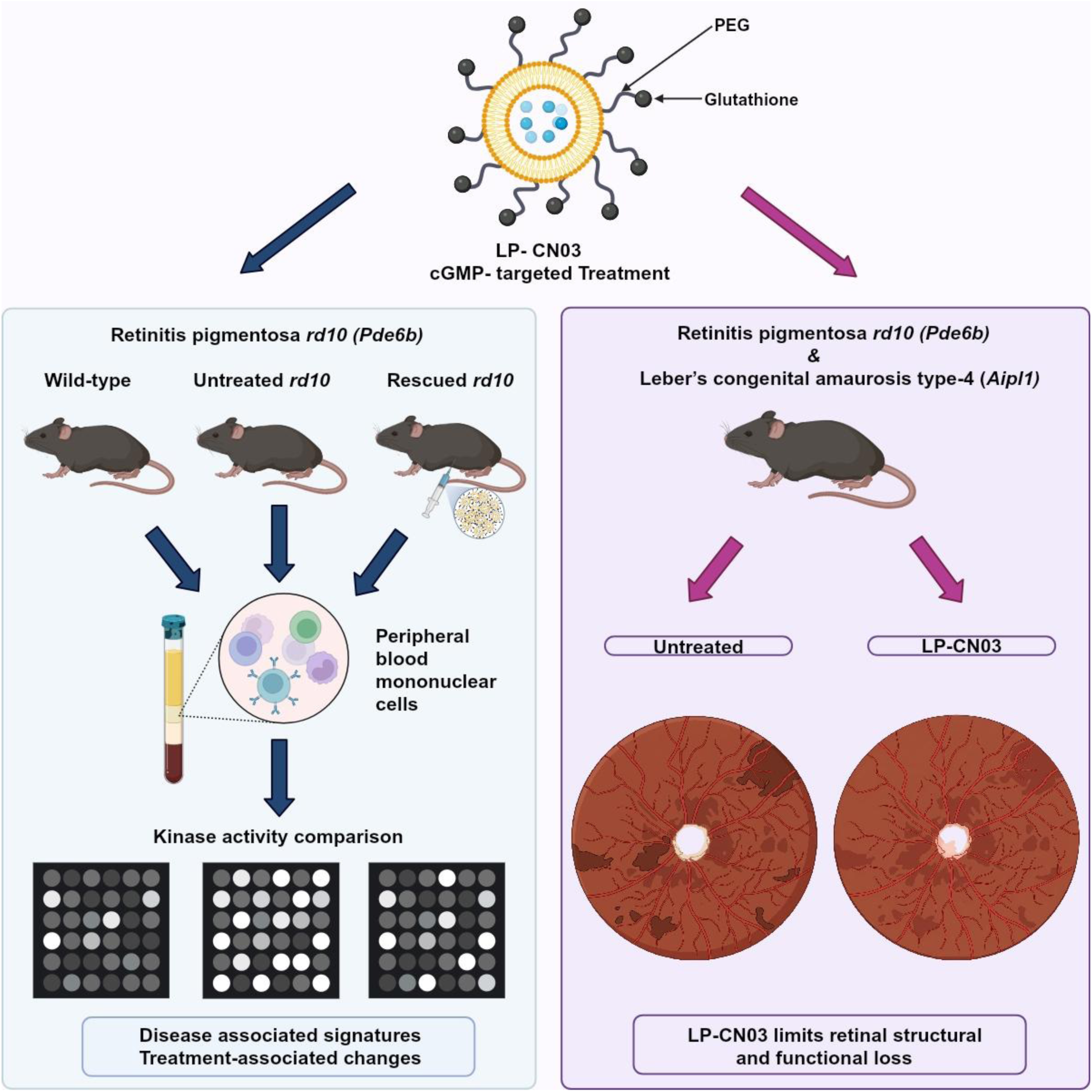

Created in BioRender. Okan, I. C. T. (2026) https://BioRender.com/76oppx6.

## Introduction

*Retinitis pigmentosa* (RP) is characterized by progressive rod and cone photoreceptor degeneration and vision loss, with different genetic causes underlying these inherited disorders. A substantial proportion of RP cases are caused by mutations affecting components of the phototransduction pathway, including phosphodiesterase 6 (PDE6). Impaired PDE6 activity causes cyclic guanosine monophosphate (cGMP) accumulation, sustained activation of cGMP-gated channels, increased calcium influx, and ultimately photoreceptor degeneration^1^. cGMP acts on cyclic nucleotide-gated channels (CNGCs) and cGMP-dependent protein kinase (PKG) to regulate photoreceptor signaling^2^. Retinal guanylate cyclases synthesize cGMP, whereas PDE6 hydrolyzes it, maintaining cGMP homeostasis^3^. Pathological cGMP accumulation drives degeneration in several inherited retinal disease models^4^. Excessive cGMP results in overactivation of PKG and CNGCs, promoting calcium overload, oxidative stress, mitochondrial dysfunction, and impaired proteostasis^4,5^. Experimental PKG inhibition markedly reduced photoreceptor cell death, suggesting that the cGMP–PKG pathway could be targeted therapeutically across genetically diverse retinal degenerations^6^.

Lebeŕs congenital amaurosis (LCA) type 4 (LCA4), caused by mutations in the *AIPL1* gene, represents a severe, early-onset retinal dystrophy, which results in severe visual function loss and blindness already during infancy or early childhood^7,8^. The photoreceptor-specific chaperone aryl hydrocarbon receptor-interacting protein-like 1 (AIPL1) supports PDE6 biosynthesis and stability by facilitating the folding and assembly of PDE6α and PDE6β subunits. Without functional AIPL1, misfolded PDE6 subunits undergo proteasomal degradation, resulting in cGMP accumulation and photoreceptor cell death^9,10^. Thus, cGMP-related RP and LCA4 share a disease mechanism that provides a common therapeutic target^6,9^. Although the PKG-targeting cGMP analog CN03 has demonstrated preclinical efficacy in animal models for RP^11^, its neuroprotective potential in LCA remains unclear.

Rp-configured cyclic nucleotide phosphorothioates were identified as stereospecific inhibitors of cyclic nucleotide-dependent protein kinases^12–14^. Early cGMP analogs had limited membrane permeability and metabolic stability, which reduced their cellular uptake and *in vivo* efficacy^15^. Subsequent lipophilic substitutions and steric modifications improved these properties and target selectivity^16–19^, leading to Rp-8-Br-PET-cGMPS (CN03)^16^. In RP models including *rd1*, *rd2*, and *rd10*, CN03 attenuated photoreceptor degeneration and preserved cone function in the *rd10* model^6,11^. The liposomal cGMP analog formulation LP-CN03 was designed to facilitate systemic delivery and retinal bioavailability. Following intraperitoneal administration, it crossed the blood–retinal barrier and preserved retinal structure and function in multiple RP mouse models, reducing rod photoreceptor loss and preventing secondary cone degeneration in the *rd10* model^11^. Based on this rationale, we evaluated systemic LP-CN03 treatment for retinal structural and functional preservation in a rapidly degenerating *Aipl1*^⁻/⁻^ mouse model of LCA4. In parallel, we characterized treatment-associated kinase signatures in peripheral blood mononuclear cells (PBMCs) from *rd10* mice to identify potential peripheral pharmacodynamic biomarkers of therapeutic response.

PBMCs are an established source of biomarkers of physiological and pathological changes^20–22^ and are used to assess dose-dependent effects and safety of drug candidates^23,24^. PBMC kinome analysis was therefore central to identifying biosurrogate markers and validating them through interventional experiments. Kinase activity analysis identified changes with potential relevance as systemic biomarkers of disease state and treatment response. Beyond the previously studied PDE6-driven RP models, LP-CN03 preserved retinal structure and function in rapidly degenerating *Aipl1*^⁻/⁻^ mice. Together, these findings suggest that cGMP-linked kinase signatures may serve as candidate diagnostic or response markers and that targeting this pathway could offer a therapeutic strategy applicable to various distinct forms of inherited retinal degeneration.

## Results

### Evaluation of LP-CN03-mediated rescue in *rd10* mice *in vivo*

Given previous evidence suggesting CN03 benefit in the *rd10* model^11^, we assessed LP-CN03 treatment in an independent cohort to confirm functional rescue using electroretinography (ERG). ERG recordings showed that photoreceptors preserved by LP-CN03 remained functionally active (Figure 1A-C). Parallel optical coherence tomography (OCT) imaging performed on 28-day-old untreated and LP-CN03–treated *rd10* mice showed a marked preservation of the photoreceptor layer in treated eyes, with structural integrity maintained across the retina, even at advanced stages of degeneration (Figure 1E). Collective analysis of retinal regions across the temporal-nasal axis revealed an overall 29.51% (40.82 ± 10.14 µm; n = 27 eyes) increased preservation of retinal layer thickness in the treated group. Droplet digital PCR (ddPCR) expression analysis (Figure 1D) revealed a significant preservation of rhodopsin (*Rho*) mRNA levels in the treated group (1.47 ± 0.34-fold; n = 16), supporting the functional and morphological benefits observed with LP-CN03 treatment. Together, the ERG, OCT, and *Rho* expression results demonstrated significant functional rescue and structural preservation by LP-CN03 in the *rd10* model. This experimental data set the base for the subsequent interrogation of peripheral PBMC markers to explore treatment-associated systemic signatures.

**Figure 1:**
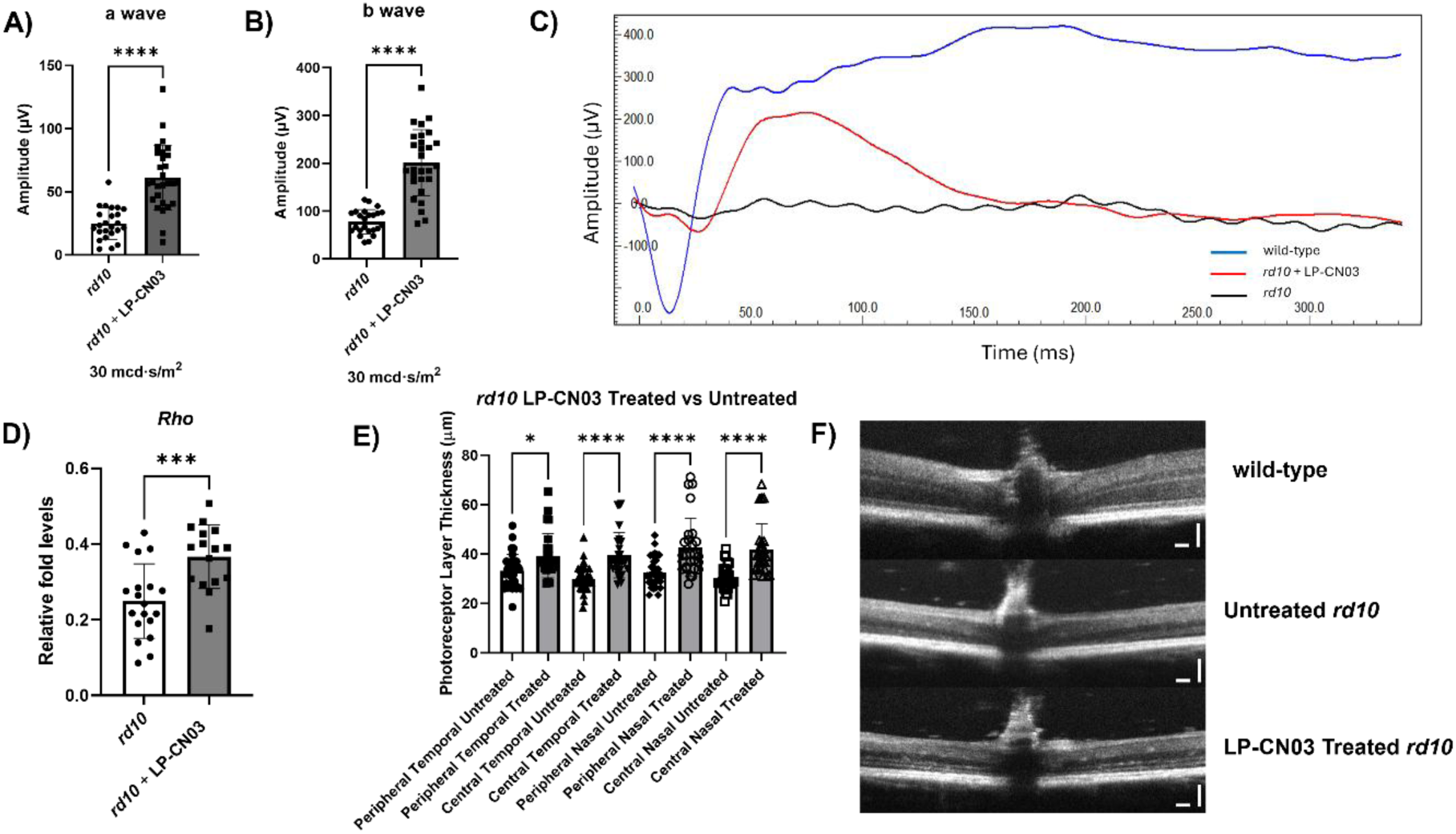
Effect of LP-CN03 treatment on retinal morphology and function in *rd10* mice. *rd10* mice were treated with an intraperitoneal dose of 30 mg/kg LP-CN03 every four days from P14 to P30. ERG recordings were performed on day 29. Electroretinography (ERG) recordings were obtained following at least 4 h of dark adaptation. Dark-adapted ERG a-wave **A**) and b-wave **B**) amplitudes from untreated and LP-CN03-treated *rd10* mice at a flash intensity of 30 mcd·s/m². **C**) Representative ERG responses (30 mcd·s/m², dark-adapted) in *rd10* animals; untreated (black), treated with LP-CN03 (red). Congenic, age-matched wild-type traces (blue) shown for comparison. Sample sizes were n = 24 eyes for untreated *rd10* and n = 28 eyes for LP-CN03-treated *rd10*. **D**) Retinal rhodopsin mRNA expression was quantified by droplet digital PCR (ddPCR) in untreated (n = 19) and LP-CN03-treated (n = 16) *rd10* mice. Retinal tissues were collected at postnatal day 30 (P30), and gene expression levels were normalized to *Gapdh* as the internal housekeeping control. **E**) Photoreceptor layer thickness in the peripheral temporal, central temporal, peripheral nasal, and central nasal retinal regions of untreated and LP-CN03-treated *rd10* mice at P28 (untreated, n = 33 eyes; LP-CN03-treated, n = 27 eyes). Measurements were obtained at distances of 948 µm from the optic nerve in the peripheral retina and 711 µm in the central retina. **F**) Representative optical coherence tomography (OCT) images from wild-type, untreated *rd10*, and LP-CN03-treated *rd10* mice at P28. Horizontal and vertical scale bars represent 100 µm. Statistical analyses were performed using GraphPad Prism software. Data are presented as mean ± SD. For comparisons involving multiple groups, ordinary one-way ANOVA followed by Šídák’s multiple-comparisons test was applied. Comparisons between two groups were performed using an unpaired two-tailed Student’s t-test. Statistical significance was defined as *p < 0.05, ***p < 0.001, and ****p < 0.0001.

### Candidate PBMC-based biosurrogate markers

A major limitation in retinal degeneration research is the absence of reliable long-term markers to assess therapeutic efficacy. Using kinome signatures, we compared disease progression between wild-type and *rd10* mice and assessed differences between untreated and LP-CN03-treated *rd10* animals. PBMC kinase activity was subsequently characterized to examine pathways associated with disease progression and treatment response. PBMCs were isolated from cardiac blood samples according to the workflow shown in Figure S1.

To demonstrate the neuroprotective effect of LP-CN03, intraperitoneal injections were administered every four days between P14 and P30, with the final dose given at P30. To standardize the pharmacokinetic effect of the compound, PBMC isolation was performed 24 hours after the last injection. Kinase activities were subsequently assessed using two distinct PamChip® peptide microarrays targeting protein tyrosine kinases (PTKs) and serine/threonine kinases (STKs), respectively. Comparison of *rd10* and wild-type mice identified 15 significantly altered phosphosites, with increases observed for 10 and decreases for 5 phosphosites in the *rd10* group (Figure 2A). Conversely, comparison of LP-CN03-treated and untreated *rd10* mice identified 16 significantly altered phosphosites, with decreases observed for 12 and increases for 4 phosphosites following treatment (Figure 2A). In the upstream kinase activity analysis, a similar pattern emerged. Comparison of *rd10* and wild-type mice showed relatively upregulated serine/threonine kinase activity, shown as red signals on the coral tree (Figure 2B). In particular, within the protein kinase A, G, and C (AGC) group, activity of cGMP-dependent PKG1 (*Prkg1*) and PKG2 (*Prkg2*) were upregulated in *rd10* mice. Some kinases among the cyclin-dependent kinases, mitogen-activated protein kinases, glycogen synthase kinases, and CDC-like kinases (CMGC) group also showed increased activity compared with controls (Figure 2B). In contrast, comparison of LP-CN03-treated and untreated *rd10* mice showed relatively decreased kinase activity, represented by blue signals on the Coral tree. Serum/glucocorticoid regulated kinase 2 (SGK2) and protein kinase C delta (PRKCD) from the AGC group, together with calcium/calmodulin-dependent protein kinase II alpha (CAMK2A) and MAP kinase-activated protein kinase 2 (MAPKAPK2) from the calcium/calmodulin-dependent protein kinase (CaMK) group, showed decreased activity following LP-CN03 treatment (Figure 2B).

**Figure 2.**
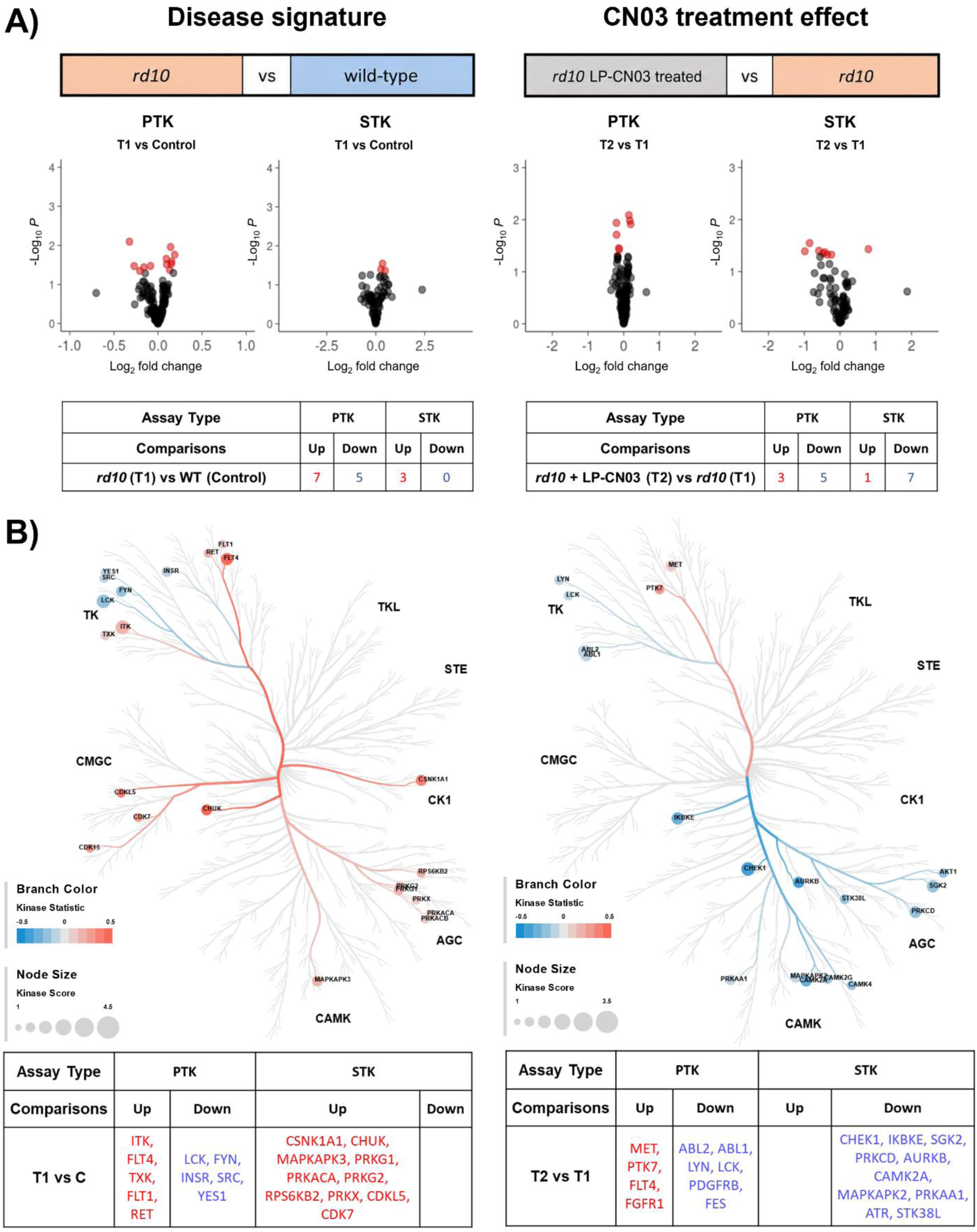
PBMC kinome profiling of *rd10* mice and its modulation by LP-CN03 treatment. **A**) Phosphosite analysis of the disease (*rd10* vs. wild-type) and treatment (LP-CN03-treated *rd10* vs. untreated *rd10*) comparisons. Wild-type control (C) mice were compared with untreated *rd10* mice (T1). T1 was compared with LP-CN03-treated *rd10* mice (T2). Volcano plots visualize statistical significance (−log₁₀ P value) versus effect size (log₂ fold change) for individual phosphosites. Red dots represent phosphosites showing significant differences (P < 0.05). Sample sizes were n = 11 for wild-type controls (C), n = 11 for untreated *rd10* mice (T1), and n = 8 (PTK) or 10 (STK) for LP-CN03-treated *rd10* mice (T2). Statistical significance was assessed using one-way ANOVA followed by Dunnett’s multiple-comparisons test. Tables summarize the numbers of significantly upregulated and downregulated phosphosites identified in the protein tyrosine kinase (PTK) and serine/threonine kinase (STK) assays. **B**) CORAL kinome trees illustrating the predicted upstream kinase activity for disease (*rd10* vs. wild-type; left) and treatment (LP-CN03-treated *rd10* vs. untreated *rd10*; right) comparisons. Only kinases with a kinase score > 1.89 are displayed. Branch color represents the predicted direction of kinase activity (red, increased; blue, decreased), whereas node size is proportional to the kinase score. Kinases are organized according to phylogenetic kinase families. The tables below list the top predicted upstream kinases identified by upstream kinase analysis in each comparison, including only kinases with kinase scores exceeding the default threshold (Kinase score > 1.3).

To gain further insight into the biological processes associated with disease state and treatment response, functional enrichment analyses were performed using the kinase signatures obtained from comparisons between *rd10* mice and wild-type controls, and between LP-CN03-treated and untreated *rd10* mice. The disease-associated kinase signature showed enrichment for receptor-mediated signaling, including vascular endothelial growth factor receptor signaling, as well as pathways regulating phosphatidylinositol 3-kinase and protein kinase B signaling (Figure 3A). The treatment-responsive signature showed enrichment for peptidyl-serine phosphorylation and pathways associated with hormone secretion and cellular signaling (Figure 3B). Integration of disease-associated and treatment-responsive kinase activity profiles further enabled the prioritization of candidate biosurrogate markers. Among these, inhibitor of nuclear factor kappa-B kinase subunit epsilon (IKBKE), CAMK2A, calcium/calmodulin-dependent protein kinase II delta (CAMK2D), SGK2, and MAPKAPK2 displayed the most consistent pattern of disease-associated activation and treatment-induced reversal, identifying them as the strongest candidate biomarkers for monitoring CN03 response (Figure 3C). A different pattern was observed for FLT4 (VEGFR3) and FGFR1, whose predicted activities increased in disease and further following LP-CN03 treatment (Figure 3C). These findings may reflect alterations in VEGF and FGF signaling, respectively, both of which have previously been implicated in RP pathogenesis and the potential activation of endogenous neuroprotective factors^25,26^.

**Figure 3.**
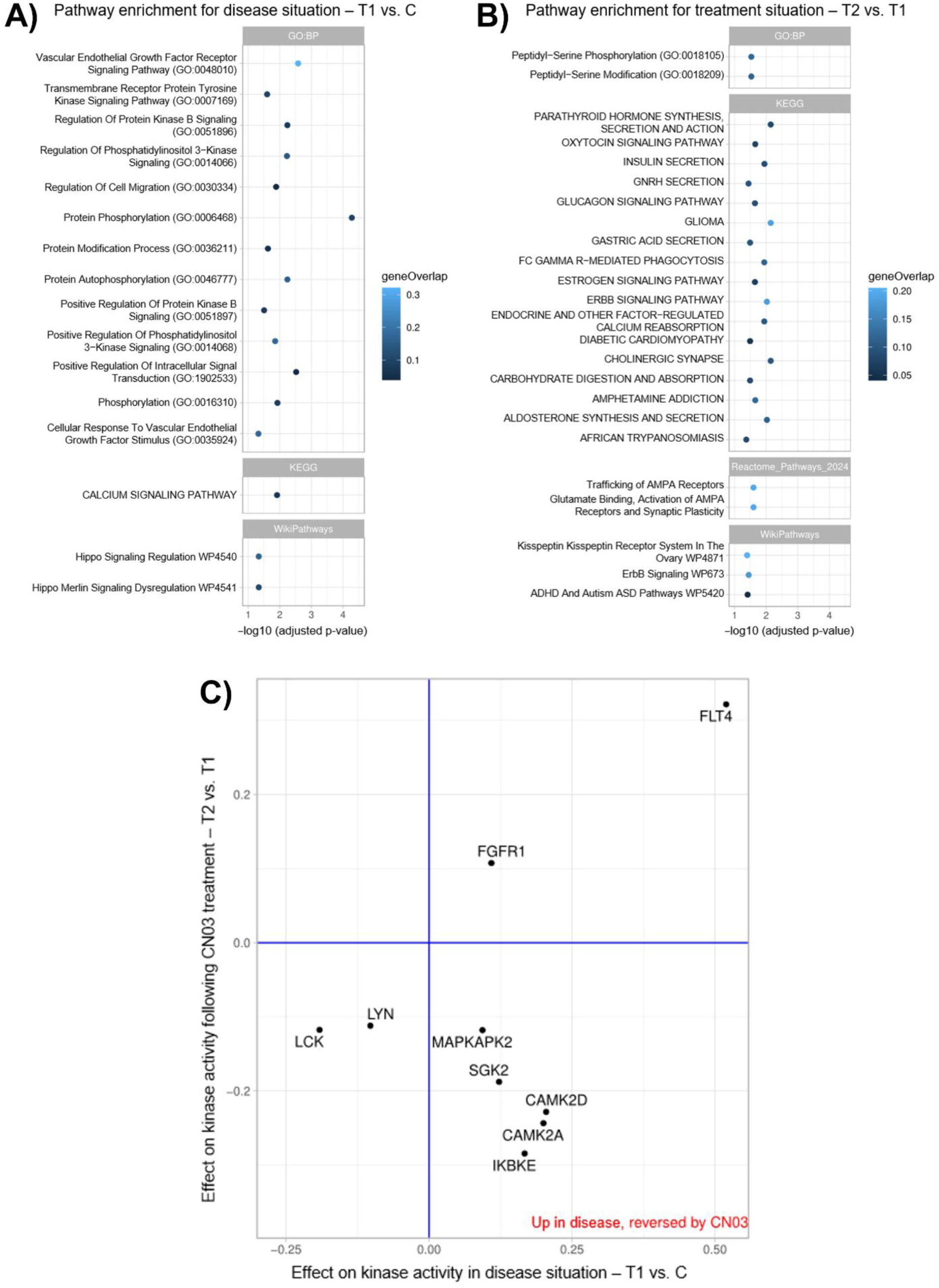
Pathway enrichment of disease-associated and CN03-responsive kinase signatures and identification of candidate biosurrogate markers. **(A)** Pathway enrichment of the disease-associated signature in untreated *rd10* mice compared with wild-type controls (T1 vs. C). (B) Pathway enrichment of the treatment-responsive signature in LP-CN03-treated *rd10* mice compared with untreated *rd10* mice (T2 vs. T1). In A and B, the horizontal axis shows −log₁₀ (adjusted p-value), and color indicates gene overlap. Analysis was performed using enrichR, with combined significant phosphosite and upstream kinase analysis (UKA) results matched by UniProt accession as input and the 338 analysed kinases/substrates as background. P values were adjusted using the Benjamini– Hochberg method, and terms with adjusted P < 0.05 were considered significant. **(C)** UKA-derived kinase activity changes in the disease comparison (T1 vs. C) plotted against changes following treatment (T2 vs. T1). IKBKE, CAMK2A, CAMK2D, SGK2, and MAPKAPK2 fall in the lower-right quadrant, indicating increased activity in disease and decreased activity following treatment, representing the most biologically relevant reversal candidates.

### LP-CN03 preserves photoreceptors in a new *Aipl1*^⁻/⁻^ LCA mouse model

To evaluate potential CN03-associated rescue effects in an LCA4 context, we created a new *Aipl1*^⁻/⁻^ mouse model. This rapidly degenerating model was generated using CRISPR/Cas9 approach to delete exon 2 and was expected to recapitulate the genetic and phenotypic consequences of the previously published ENU-derived *Aipl1*^tvrm119^ mutation (MMRRC stock no. 043590-JAX). The induced c.276+5G>A mutation was predicted to cause exon 2 skipping through a splicing defect ^27^. Exon 2 encodes part of the FK506-binding protein-like (FKBP-like) domain involved in AIPL1 chaperone function and PDE6 stabilization. Its deletion removes 60 amino acids (residues 33–92) in frame, eliminating much of this domain and thereby disrupting AIPL1 function. This loss of function results in severe early-onset retinal degeneration. The genomic organization of the *Aipl1* locus and the CRISPR/Cas9 strategy used to delete exon 2, as well as the editing strategy and the single-guide RNA (sgRNA) target sites are shown in Figure 4A. PCR-based genotyping and Sanger sequencing confirmed the 203 bp deletion (Figure 4B-C). Following colony generation, *Aipl1*^⁻/⁻^ mice were bred to homozygosity and subsequently characterized to evaluate retinal degeneration and therapeutic response.

**Figure 4.**
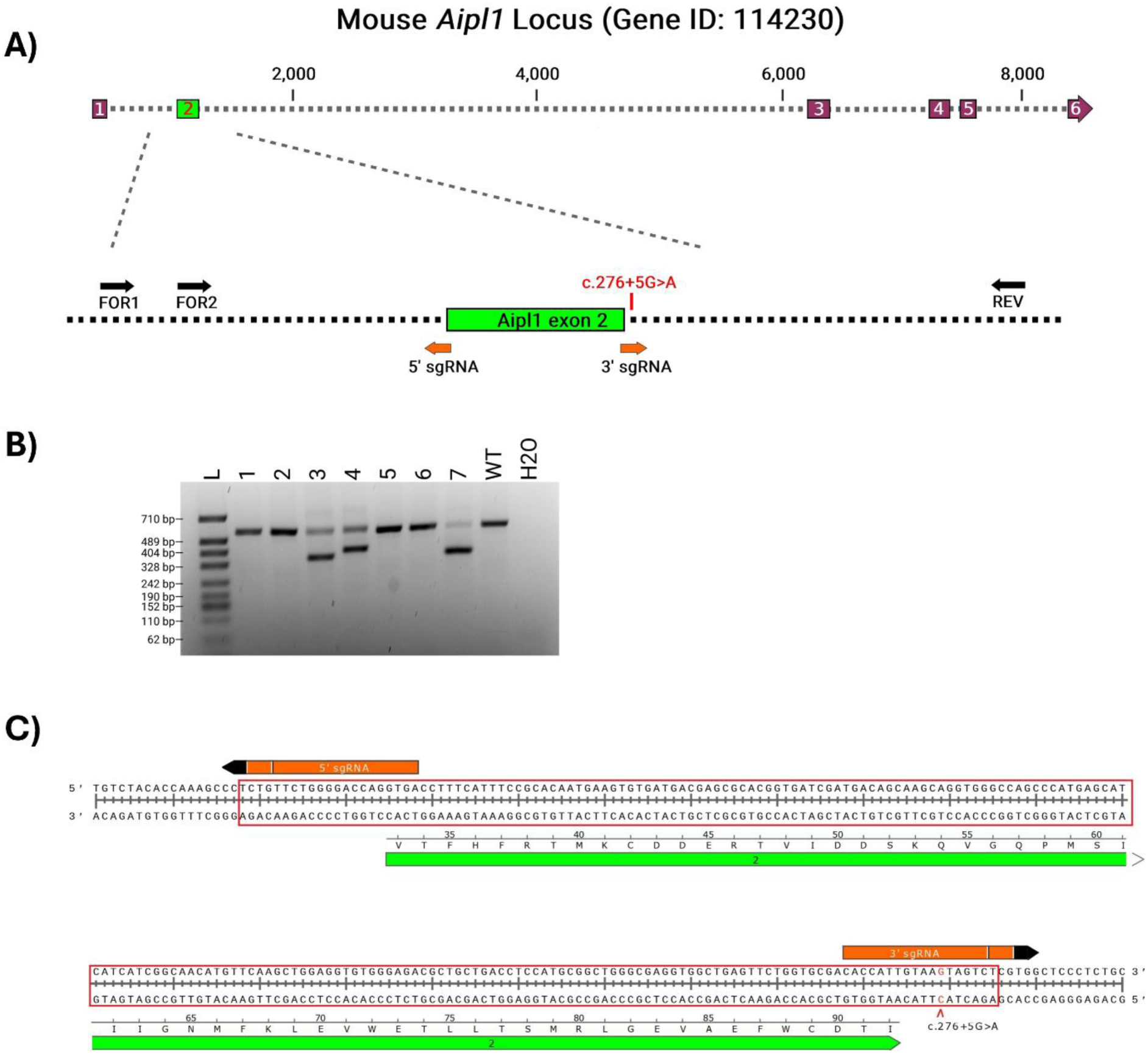
**Generation of the *Aipl1*^⁻/⁻^ mice using CRISPR–CasG editing**. **A**) Schematic representation of the mouse *Aipl1* genomic locus and enlargement of the region surrounding exon 2. The positions of the 5′ and 3′ single-guide RNAs (sgRNAs) flanking exon 2, the c.276+5G>A splice-site variant in *Aipl1^tvrm11S^* mice, and the primer binding sites (FOR1, FOR2 and REV) used for PCR genotyping and sequencing are indicated. The primer pairs were designed to detect the exon 2 deletion based on PCR amplicon size differences. **B**) Agarose gel showing PCR-based genotyping of founder mice born following embryo transfer. Lanes 1–7 represent founder mice; WT, wild-type control; H₂O, no-template control; L, DNA size ladder. **C**) Sequence illustration of the CRISPR–Cas9 editing strategy used to generate the *Aipl1*^⁻/⁻^ mice. The 5′ and 3′ sgRNAs were designed to excise exon 2, resulting in deletion of the membrane-binding domain, thereby generating a nonfunctional AIPL1 protein. Red arrowhead indicates position of the c.276+5G>A splice-site variant. Red rectangles denote deleted sequence in the *Aipl1* KO allele.

For characterization of the model, two months and six months old animals were first evaluated by fundus imaging and OCT (Figure 5). Consistent with previous descriptions, our *Aipl1*^⁻/⁻^mice displayed a grainy fundus appearance with attenuated retinal vessels that progressed to a patchy retinal phenotype, previously associated with glial scarring and advanced photoreceptor degeneration^27^. OCT imaging revealed a marked reduction in photoreceptor layer thickness, consistent with advanced retinal degeneration. As a tangible proxy for photoreceptor layer thickness, in OCT images the retinal thickness was measured from the inner boundary of the inner nuclear layer (INL) to the outer boundary of the photoreceptor outer segments (OS), encompassing the INL, outer plexiform layer (OPL), outer nuclear layer (ONL), external limiting membrane (ELM), photoreceptor inner segments (IS), and photoreceptor outer segments (OS). Measurements were obtained from four retinal locations corresponding to the temporal peripheral, temporal central, nasal peripheral, and nasal central regions. When wild-type control values were normalized to 100%, the mean photoreceptor layer thickness across these four regions was reduced to 24.3% in 2-month-old

**Figure 5.**
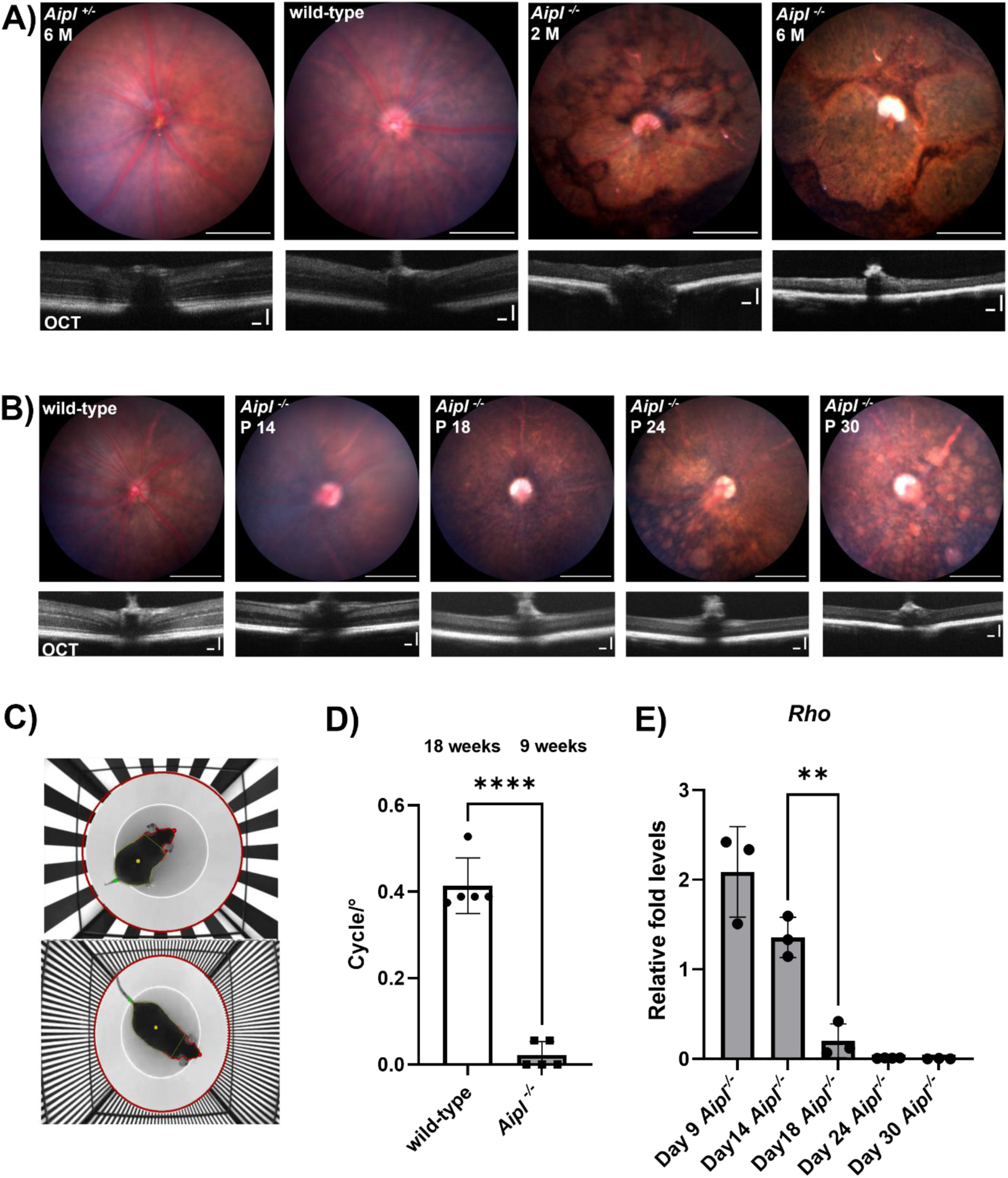
Characterization of the newly created *Aipl1*^⁻/⁻^ mice. **A**) Time dependent evaluation of retinal degeneration in *Aipl1^-/-^*mice using fundus and optical coherence tomography (OCT) imaging. Representative images from *Aipl1^-/-^* mice at 2 and 6 months of age are shown. Control images include 6-month-old heterozygous and wild-type mice. **B**) Longitudinal monitoring of retinal structure in *Aipl1^-/-^* mice. Fundus and OCT images were collected at postnatal days 14, 18, 24, and 30. Fundus imaging revealed attenuation of retinal vessels and the development of a patchy retinal appearance, consistent with progressive retinal degeneration, while OCT demonstrates progressive thinning and degeneration of the photoreceptor layer. For **(A)** and **(B)**, scale bars represent 500 µm in fundus images and 100 µm in OCT images in both the horizontal and vertical directions. **C**) Visual function impairments in *Aipl1^-/-^* mice measured using the optomotor response (OMR). Representative screenshots of OMR tracking at different spatial frequencies are shown. **D**) Quantification of spatial frequency thresholds (cycles/°) in 18-week-old wild-type control mice (n = 5) and 9-week-old *Aipl1^-/-^* mice (n = 5). Automated analysis showed a marked reduction in visual performance in *Aipl1^-/-^* mice; controls detected an average spatial frequency of 0.414 cycles/°, whereas knockouts reached only 0.022 cycles/°. Data are presented as mean ± SD. Statistical significance was assessed using two-tailed, unpaired Student’s t-tests (**** p < 0.0001). **E**) Retinal rhodopsin (*Rho*) mRNA expression in *Aipl1^-/-^* mice at postnatal days 9, 14, 18, 24, and 30 was quantified using droplet digital PCR (ddPCR) and normalized to *Gapdh*. Each data point represents an individual animal (n = 3 per time point, except P24, n = 4), and bars represent mean ± SD. Statistical significance was assessed using Welch’s one-way ANOVA, followed by Dunnett’s T3 multiple-comparisons test for comparisons between consecutive time points. **P < 0.01.

*Aipl1*^⁻/⁻^ mice, corresponding to an approximately 75.7% decrease relative to controls. Degeneration progressed further by 6 months of age. Mean photoreceptor layer thickness declined to 13.8%, corresponding to an approximately 86.2% reduction. Figure 5A shows representative fundus and OCT images from *Aipl1*^⁻/⁻^ and control mice at 2 and 6 months. Control images were obtained from 6-month-old heterozygous and wild-type mice.

To assess behavioral visual function in the CRISPR-generated *Aipl1*^⁻/⁻^ mice, optomotor response (OMR) was assessed (Figure 5C, 5D). Whereas wild-type controls demonstrated spatial-frequency thresholds around 0.417 cycles/*°*, *Aipl1*^⁻/⁻^ mice had difficulty responding even at the initial spatial frequency tested, indicating a profound loss of reflex-based visual function. These results provided functional support for the severe degeneration documented by OCT and fundus imaging. To determine the onset of retinal degeneration, longitudinal fundus and OCT imaging were performed. Longitudinal fundus imaging demonstrated progressive disease-associated changes, characterized by the gradual development of a granular retinal appearance, attenuation of retinal vessels, and the appearance of patchy retinal lesions at later time points. Because RP progresses later in life, whereas LCA4 is marked by rapid degeneration beginning in early childhood, a similarly accelerated pattern was anticipated in *Aipl1*^⁻/⁻^ mice. OCT scans at P14, *i.e.* just after eye opening, already revealed substantial photoreceptor damage, particularly within the ONL (Figure 5B). Degeneration advanced rapidly at P18 and P24, with only a thin residual photoreceptor layer remaining by P30. Representative haematoxylin and eosin (HCE) images comparing age-matched wild-type and *Aipl1*^⁻/⁻^ retinas, illustrated the progressive loss of the photoreceptor layer in the *Aipl1*^⁻/⁻^ mice (Figure S4). Quantitative OCT measurements further demonstrated the progressive reduction in photoreceptor layer thickness between P14 and P30 (Figure S5). Consistent with these structural findings, ddPCR analysis showed a continuous decline in *Rho* expression, with significant reductions between P14 and P18 (Figure 5E).

After characterization of the new *Aipl1*^⁻/⁻^ mice, we wanted to assess whether CN03 could exert neuroprotective effects also in LCA4. Due to the severe and early-onset degeneration observed in this model, CN03 treatment was initiated earlier with an increased dosing frequency, following the strategy used in *rd10* animals. *Aipl1*^⁻/⁻^ mice therefore received intraperitoneal injections of the liposomal CN03 formulation (LP-CN03; 30 mg/kg) on postnatal days 7, 10, 12, 14, 17, 19, and 21.

Representative fundus and OCT images acquired before and after treatment at P18 and P23 are shown in Figure 6A. At day 23, LP-CN03-treated *Aipl1*^⁻/⁻^ mice showed qualitatively better preservation of fundus appearance, characterized by a more homogeneous retinal background and fewer and less extensive multifocal pale fundus abnormalities compared with untreated controls (Figure S2). Photoreceptor layer thickness measurements from P18 OCT scans show a significantly greater preservation of the photoreceptor layer in LP-CN03–treated mice compared to untreated controls, particularly at the central temporal and peripheral positions (Figure 6B). Due to the very rapid *Aipl1* mutation-induced degeneration, by P23, retinal thickness measurements showed no statistically significant differences between treated and untreated animals (Figure 6B). Representative HCE images illustrate an improved preservation of the photoreceptor layer in CN03-treated *Aipl1*^⁻/⁻^ mice compared with untreated controls (Figure S3).

**Figure 6.**
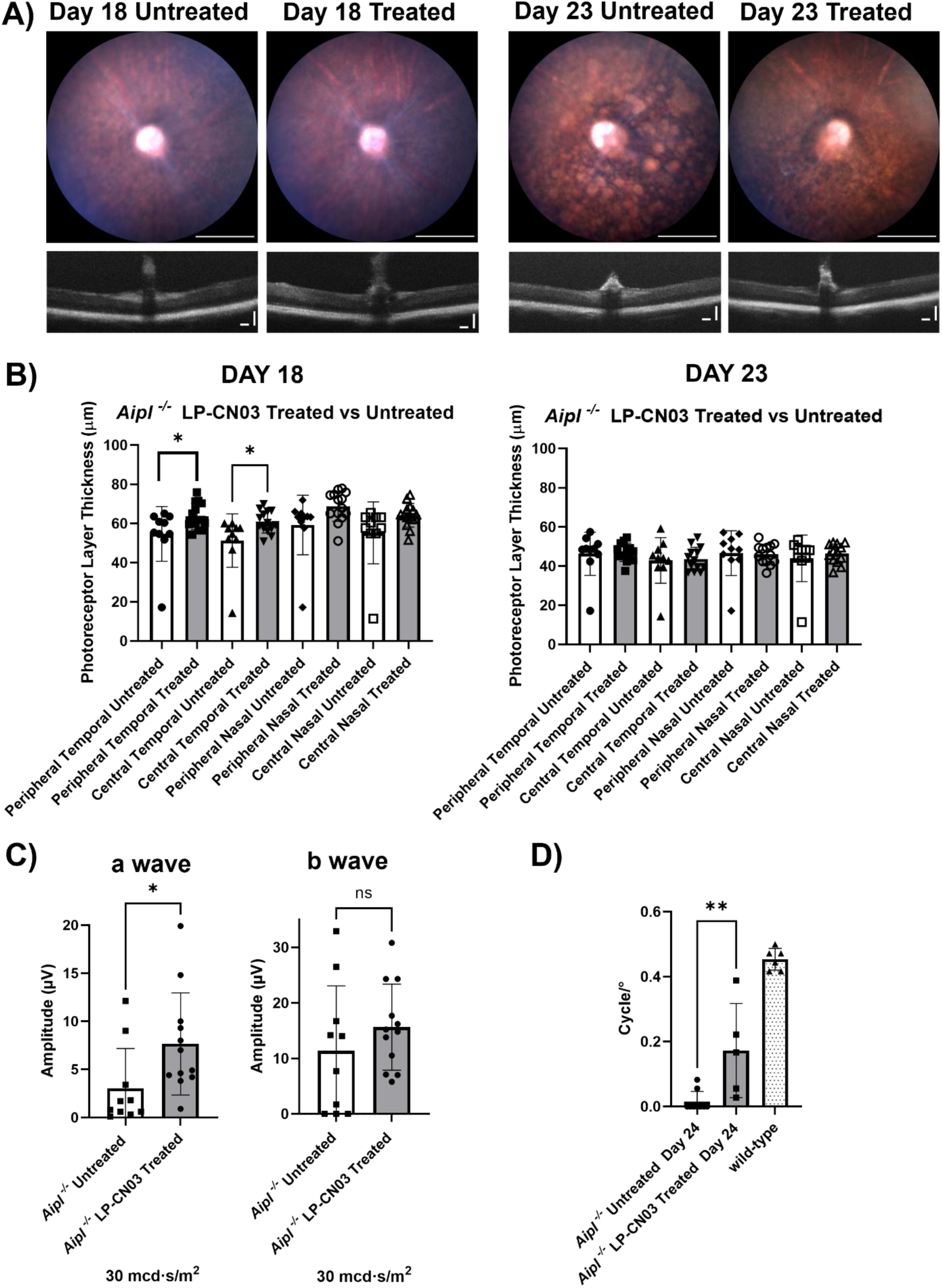
Therapeutic effects of LP-CN03 in *Aipl1^-/-^* mice. **A**) Representative fundus and optical coherence tomography (OCT) images of *Aipl1^-/-^* mice treated with LP-CN03. Images show eyes of untreated and treated mice at post-natal day (P) 18 (left) and 23 (right). Scale bars represent 500 µm in fundus images and 100 µm in OCT images in both the horizontal and vertical directions. Note the marked differences in retinal structure and morphology between treated and untreated *Aipl1^-/-^* mice. **B)** Quantification of retinal layer thickness via OCT in LP-CN03-treated and untreated *Aipl1^-/-^* mice at P18 (treated, n = 14 eyes; untreated, n = 9 eyes) and P23 (treated, n = 13 eyes; untreated, n = 9 eyes). Retinal thickness was measured in central and peripheral retina, in the temporal and nasal quadrants. Data are presented as mean ± SD. Statistical comparisons were performed using two-tailed, unpaired Student’s t-tests in GraphPad Prism, with p < 0.05 considered significant. **C**) Scotopic electroretinography (ERG) evaluation of retinal function in LP-CN03–treated versus untreated *Aipl1^-/-^* mice. ERG recordings were conducted at P24 under dark-adapted conditions using a 30 mcd·s/m² light stimulus. Group sizes were n = 12 eyes for treated and n = 10 eyes for untreated mice. Data are presented as mean ± SD. Statistical analysis was performed using two-tailed, unpaired Student’s t-tests in GraphPad Prism, with p < 0.05 indicating significance (*). **D**) Optomotor response analysis following LP-CN03 treatment. At P24, visual acuity was assessed to determine the neuroprotective effect of LP-CN03. Untreated *Aipl1^-/-^* mice displayed markedly reduced spatial frequency thresholds (mean = 0.01544 cycles/°), whereas LP-CN03–treated mice showed significant improvement (mean = 0.1724 cycles/°). Age matched wild-type control mice exhibited a mean response of 0.4537 cycles/°. Group sizes were: untreated knockout (n = 9), treated (n = 5), and wild-type controls (n = 6). Data are presented as mean ± SD. Statistical comparisons were performed using two-tailed, unpaired Student’s t-tests in GraphPad Prism. **p < 0.01.

ERG and OMR assessments provided additional evidence of the neuroprotective effect of LP-CN03 showing improvements in both retinal function and visual performance. Scotopic ERG recordings were obtained at P24 after a minimum of four hours of dark adaptation to assess retinal function. At a stimulus intensity of 30 mcd·s/m², LP-CN03–treated *Aipl1*^⁻/⁻^ mice showed a significant increase in a-wave amplitude compared with untreated animals, whereas b-wave amplitudes did not differ significantly between groups (Figure 6C). Because the a-wave reflects the electrical activity of photoreceptors, particularly in rod cells under scotopic conditions^28^, recovery in treated animals indicates improved photoreceptor function and suggests that LP-CN03 provided protective effects at the outer retinal level. Because degeneration in the *Aipl1*^⁻/⁻^ mice progresses extremely rapidly, only limited photoreceptor survival was expected. Therefore, we included OMR analysis to determine whether this limited structural preservation translated into measurable visual function. In the untreated group (n = 9), seven animals failed to pass the initial step of the OMR test and therefore received a score of zero. LP-CN03–treated *Aipl1*^⁻/⁻^ mice showed a markedly better optomotor response than untreated animals. On P24, untreated mice displayed an average threshold of 0.01544 cycles/°, whereas treated mice achieved a mean spatial frequency of 0.1724 cycles/° (Figure 6D).

Retinas from LP-CN03-treated and untreated mice were analyzed by ddPCR for photoreceptor-associated gene expression at P24. *Rho* expression was markedly higher in LP-CN03-treated mice compared with untreated controls, indicating preservation of *Rho* levels following LP-CN03 treatment (Figure 7A). ddPCR analysis revealed a significant increase in the cone opsin *Opn1mw* (Figure 7B) and the rod-specific *Gnat1* transcript levels (Figure 7D) in treated *Aipl1*^⁻/⁻^ mice compared with untreated littermates. In contrast, *Opn1sw* (Figure 7C) and *Gnat2* (Figure 7E) showed modest increases in treated animals, but these differences did not attain statistical significance. These findings indicate improved preservation of rod phototransduction components, as reflected by increased *Gnat1* expression. The significant increase in *Opn1mw* suggests partial preservation of M-cones, whereas the absence of significant changes in the S-cone opsin *Opn1sw* and cone-specific *Gnat2* indicates that cone phototransduction components were not substantially restored under these conditions. Even at an advanced stage of *Aipl1*^⁻/⁻^ photoreceptor degeneration, both the OMR assessments and ddPCR data demonstrated a partial preservation following LP-CN03 treatment.

**Figure 7:**
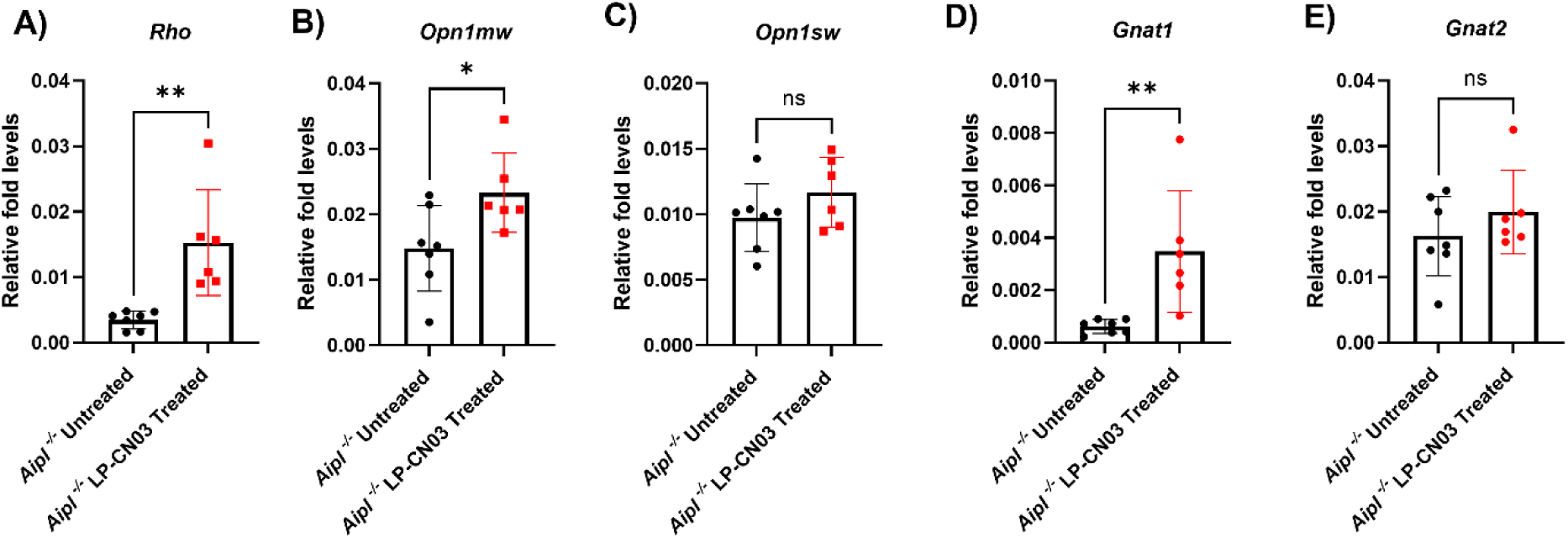
ddPCR analysis of gene expression after LP-CN03 treatment. Droplet digital PCR (ddPCR) was used to quantify retinal mRNA levels of **A**) rhodopsin (*Rho*), **B**) *Opn1mw* (M-cone opsin), **C)** *Opn1sw* (S-cone opsin), **D**) *Gnat1*, and **E**) *Gnat2* in *Aipl1^-/-^* mice that were either untreated (n = 7) or treated with LP-CN03 (n = 6). Retinas were collected at P24. Gene expression levels were normalized to *Gapdh*, which served as reference housekeeping gene. Data are presented as mean ± SD. Statistical significance was assessed using two-tailed, unpaired Student’s t-tests in GraphPad Prism. *P < 0.05; **P < 0.01

## Discussion

Inherited retinal degenerations, such as RP and LCA, cause progressive vision loss, while their genetic heterogeneity and limited systemic biomarkers complicate disease characterization and monitoring.

Here, we show that treatment with LP-CN03 preserved retinal structure and function in RP and LCA models *in vivo*, highlighting cGMP-signaling as a common target for genetically distinct retinal diseases. Moreover, kinome profiling of PBMCs distinguished healthy from diseased states and showed treatment efficacy in *rd10* mice, demonstrating its potential as a translational biomarker.

### Therapeutic implications of LP-CN03 in inherited retinal degeneration

Previous studies showed that pharmacological inhibition of PKG using systemic LP-CN03 treatment reduced photoreceptor cell death in the *rd1*, *rd2*, and *rd10* mouse models for RP, with functional rescue demonstrated in *rd2* and *rd10* mice^6,11^. In the present study, we independently confirmed the neuroprotective effect of LP-CN03 in the *rd10* model and proceeded with an evaluation of therapeutic efficacy in a newly generated mouse model of *AIPL1*-associated LCA4. Compared with the more slowly progressing *rd10* model, *Aipl1*^⁻/⁻^ induced degeneration in the mouse retina began before eye opening, requiring treatment initiation at P7. Despite this narrow therapeutic window and the rapid disease progression, LP-CN03 preserved retinal morphology, photoreceptor-specific gene expression, and retinal function. Systemic LP-CN03 treatment was therefore effective in a genetically distinct, and very severe model of inherited retinal degeneration. Systemic administration was chosen to avoid repeated intraocular injections and the associated cumulative risk of endophthalmitis ^29^. Because retinal drug delivery is limited by the blood–retinal barrier^30^, CN03 was encapsulated in glutathione-targeted PEGylated (GSH-PEG) liposomes, a formulation previously shown to prolong systemic circulation and facilitate transport across barrier tissues^11,31^. The preservation of retinal morphology and function observed in this study provided additional evidence for the feasibility of this delivery strategy, including in the rapidly degenerating *Aipl1*^⁻/⁻^ mice. The formulation itself is another important consideration for optimizing PKG-targeted therapy. A recent pharmacokinetic study in rabbits demonstrated that PEGylated liposomal formulations had longer intraocular retention than free CN03, while lipid composition further influenced drug release kinetics^32^. These observations indicate that optimization of liposomal composition could further extend intraocular drug exposure.

The encouraging efficacy of LP-CN03 observed across multiple inherited retinal degeneration models has also stimulated the development of next-generation PKG inhibitors. CN238 represents a second-generation cGMP analog derived from CN03 that retains the same mechanism of PKG inhibition while exhibiting improved inhibitory potency. CN238 was reported to preserve photoreceptor viability and function in *rd1*, *rd10*, and *Rho*^I255del^ retina and to protect retinal ganglion cells in axotomized retinal explants^33,34^. Together with the present findings, these studies further support PKG inhibition as a promising therapeutic strategy for gene-and mutation-independent treatment of inherited retinal degeneration.

While PKG-targeted neuroprotection represents a mutation-independent therapeutic strategy, gene replacement therapy has recently emerged as another promising approach for *AIPL1*-associated retinal degeneration. In a first-in-human clinical study, four children with biallelic disease-causing variants in *AIPL1* were treated with the AAV-based gene replacement vector rAAV8.hRKp.AIPL1, which carries the human *AIPL1* coding sequence^35^. Subretinal delivery of this vector has been shown to provide sustained improvements in visual function and preservation of outer retinal structure in young children with severe *AIPL1*-associated retinal dystrophy. In this therapeutic landscape, strategies targeting downstream degenerative pathways may serve as supportive approaches that complement mutation-specific gene replacement therapies.

### PBMC kinase signatures as pharmacodynamic biomarkers of treatment response

Assessing treatment response in RP is challenging because of the slow course of the disease and structural changes in photoreceptors that are difficult to detect at early stages. Here, biosurrogate markers could provide a much-needed, more immediate measure of therapeutic efficacy. A central aim of this work was to examine changes in kinase signatures associated with photoreceptor degeneration and their potential use as measurable indicators of disease progression. For this purpose, kinase profiling was performed using PBMCs, which provide an accessible and minimally invasive source for pharmacodynamic biomarker discovery. Disease-associated alterations in kinase activity were characterized in the *rd10* model as a basis for identifying biomarkers responsive to treatment. Increased activity detected within the AGC kinase family, and particularly PKG1 and PKG2, was consistent with excessive cGMP-dependent signaling, which directly drives photoreceptor cell death. This is supported by earlier findings on reduced photoreceptor cell death after pharmacological inhibition of PKG^6,11^.

A second important objective was the identification of accessible pharmacodynamic biomarkers of treatment response based on the demonstrated therapeutic effects of LP-CN03 in inherited retinal degeneration. Previous work has suggested that PKG inhibition induces overlapping kinase responses in retinal tissue and PBMCs, supporting the concept that peripheral kinase activity can reflect disease-associated signaling changes^36^. This approach enabled the identification of treatment-responsive candidate biosurrogate kinases, including IKBKE, CAMK2A, CAMK2D, SGK2, and MAPKAPK2. All five candidates displayed increased activity in *rd10* PBMCs relative to wild-type controls and a reversal of this pattern following LP-CN03 treatment, which supports their potential as pharmacodynamic markers of treatment response. To our knowledge, this is the first study to perform biomarker profiling in a treated RP model. Remarkably, several of the treatment-responsive kinases identified in the present study have also been reported following PKG inhibition in other studies:

An *ex vivo rd10* retinal explant study reported decreased CaMK family activity following CN03 treatment^37^. Complementary phosphoproteomic analysis in the same study further identified modulation of MAPK-associated signaling. Similarly, a recent rabbit study found reduced activity of CaMK family kinases, IKBKE, and AMP-activated protein kinase catalytic subunit α1 (*PRKAA1*) in PBMCs following CN03 treatment^36^. The identified candidate kinases participate in several signaling pathways involved in calcium signaling, cellular stress responses, immune regulation, and cellular homeostasis. CAMK2A and CAMK2D are Ca²⁺/calmodulin-dependent kinases that regulate calcium-dependent neuronal signaling^38^, whereas MAPKAPK2 is a downstream effector of the p38 MAPK pathway that regulates cellular stress and inflammatory signaling^39^. IKBKE is a non-canonical IκB kinase that regulates NF-κB-associated signaling^40^, and SGK2 belongs to the serum/glucocorticoid-regulated kinase family involved in the regulation of ion transport and cellular homeostasis^41^. CaMK2 activity has previously been implicated in retinal signaling and photoreceptor degeneration^42,43^. Similarly, changes in MAPK signaling have been linked to retinal stress responses^44^. Beyond these functional associations, the relevance of the identified candidates as peripheral biomarkers is supported by overlapping treatment-responsive signaling across retinal and peripheral compartments. CN03-responsive changes in CaMK-and MAPK-associated signaling have previously been observed in *rd10* retinal explants^37^, while modulation of CaMK family kinases and IKBKE was independently detected in PBMCs following CN03 treatment^36^. Together with their disease-associated increase and reversal following LP-CN03 treatment in the present study, these findings suggest that the identified kinase panel may reflect systemic signaling changes associated with retinal treatment response.

Overall, this study confirms the previously reported neuroprotective effects of LP-CN03 in the *rd10* model and extends these findings to the rapidly degenerating *Aipl1*^⁻/⁻^ mice. These effects are supported by converging structural and functional evidence. The translational relevance of these findings is further strengthened by systemic PBMC profiling, which may constitute the first application of kinome analysis in rescued animals, identifying signaling markers associated with therapeutic response. Together, these findings support LP-CN03 as a gene-agnostic neuroprotective strategy and show the translational potential of PBMC kinome profiling as a pharmacodynamic biomarker platform for inherited retinal degeneration.

## Materials and Methods

### Animals

The *Aipl1*^⁻/⁻^ mice used in this study were generated on a C57BL/6J background using a CRISPR/Cas9-based strategy. C57BL/6JRj control mice were obtained from Janvier Labs, whereas *rd10* mice were obtained from The Jackson Laboratory (strain #004297). This study was approved by the Animal Experiments Ethics Committee of Acıbadem Mehmet Ali Aydınlar University and carried out in accordance with the ARVO Statement for the Use of Animals in Ophthalmic and Vision Research. All mice were maintained at 22–24°C under a 12-hour light/dark cycle, with *ad libitum* access to water and standard mouse chow.

### Generation of *Aipl1*^⁻/⁻^ mice

*Aipl1*^⁻/⁻^ mice were developed by CRISPR/Cas9 based genome editing of mouse zygotes. Two single guide RNAs (sgRNAs) (5′-TCACCTGGTCCCCAGAACAG-3′ and 5′-CACCATTGTAAGTAGTCTCG-3′) targeting regions flanking the second coding exon of the mouse *Aipl1* gene (Gene ID: 114230; MGI: 2148800) were selected to delete the flanked sequence. sgRNAs were incubated with Cas9 protein to form Cas9/sgRNA complexes, which were subsequently electroporated to embryos. Embryos were transferred to pseudopregnant CD1 females to generate founders. Previously established protocols were utilized for CRISPR gene targeting/genotyping strategy design, handling of animals, and manipulation of embryos ^45,46^.

Genotyping of newborn pups was performed using tail biopsies prepared as previously described ^47^. PCR genotyping was performed using primers FOR1 and REV (Table S2), which amplify a 563-bp fragment spanning exon 2 of the wild-type *Aipl1* allele. In CRISPR-edited animals, deletion of 203 bp results in a shorter knockout amplicon of 360 bp. Correctly edited alleles were confirmed by Sanger sequencing of the deletion allele using FOR2 primer.

### Preparation and administration of LP-CN03

*Aipl1*^⁻/⁻^ mice and *rd10* mice received intraperitoneal injections of glutathione-targeted PEGylated liposomal CN03 (GSH-PEG LP-CN03). GSH-conjugated liposomes obtained from Mireca Medicines (Tübingen, Germany) were used to encapsulate CN03 using a multi-step formulation method ^11^. CN03 was remotely loaded into a Ca²⁺ acetate–based lipophilic carrier for a final concentration of 5 mg/mL. CN03 (sodium salt) was dissolved in sterile water, and 500 μL of this solution was added to the Iyo Lipo CaAc formulation. The mixture was incubated at 60 °C for 45 minutes with gentle agitation every 15 minutes and stored at 4 °C until use. Prior to each injection, animals were weighed and the injection volume was adjusted to deliver 30 mg/kg LP-CN03.

### Fundus and optical coherence tomography (OCT) imaging and quantification

Fundus and optical coherence tomography (OCT) imaging were performed under isoflurane anesthesia following pupillary dilation with a topical ophthalmic solution containing 0.5% tropicamide and 5% phenylephrine hydrochloride. Fundus images were acquired using the MICRON IV retinal imaging system (Phoenix-Micron, Inc., Bend, OR, USA), whereas OCT images were obtained using the iVivo OCT system (OcuScience, Henderson, NV, USA). A 0.2% carbomer gel (Viscotears®, Bausch + Lomb, Kingston upon Thames, UK) was applied to maintain corneal hydration. For retinal layer analysis, volumetric OCT imaging covered the region surrounding the optic nerve head. Frames with stable image quality were selected and imported into ImageJ as a stack. Z-projection of the image stack improved resolution and reduced background signal. Retinal layer thickness was measured manually in the temporal and nasal regions, at distances of 150 pixels (711 µm) and 200 pixels (948 µm) from the optic nerve head. Eyes with OCT images with low resolution for quantification were excluded from the analysis. The number of eyes included in each analysis is indicated in the corresponding figure.

### Electroretinography (ERG)

Retinal electrophysiology was evaluated using the HMsERG LAB System (OcuScience, Henderson, NV, USA). Animals were dark-adapted for at least 4 hours before scotopic ERG recording, and baseline electrical noise was checked before data acquisition. Mice were anesthetized with isoflurane, and pupils were dilated with a topical ophthalmic solution containing 0.5% tropicamide and 5% phenylephrine hydrochloride. A 0.3% hypromellose ophthalmic gel (GenTeal® Tears Severe Lubricant Eye Gel, Alcon Laboratories, Inc., Fort Worth, TX, USA) was applied to the cornea for hydration and contact with the recording electrode. The scotopic ERG parameters are provided in Table S3. ERG traces were processed with the manufacturer’s software using a 20 Hz low-pass filter, a 50 Hz notch filter, and a spike filter to reduce background electrical noise. The system’s integrated analysis tools were used to determine a-and b-wave amplitudes from the processed waveforms. ERG recordings with waveform distortion that prevented reliable amplitude measurements were excluded. Sample sizes are reported in the corresponding figures.

### Optomotor response (OMR)

Visual function was assessed using the OptoDrum system designed for automated measurements (Striatech, Tübingen, Germany). Mice were tested without anesthesia and placed individually on the central platform. Visual acuity was measured by recording optomotor tracking responses to rotating vertical black and white gratings of progressively increasing spatial frequency. The OptoDrum software was used to automatically track and analyze head movements.

### Retinal tissue preparation and cryosectioning

Whole eyes were fixed in 4% paraformaldehyde (PFA; Sigma-Aldrich, St. Louis, MO, USA; Cat. No. 158127) for 1 h at room temperature and subsequently washed with Dulbecco’s phosphate-buffered saline without calcium and magnesium (DPBS; Gibco, Thermo Fisher Scientific, Waltham, MA, USA; Cat. No. 14190-094) at 4 °C. A small corneal incision was then made to facilitate reagent penetration, followed by post-fixation in 4% PFA for 2 h at 4 °C. The eyes were washed with DPBS at 4 °C and incubated overnight in 30% sucrose (Duchefa Biochemie, Haarlem, the Netherlands; Cat. No. S0809) at 4 °C. Following sucrose infiltration, the samples were washed with DPBS, embedded in Tissue-Tek O.C.T. compound (Sakura Finetek, Torrance, CA, USA), and rapidly frozen using liquid nitrogen. Cryosections were obtained at a thickness of 10 μm using a CryoStar NX50 cryostat (Thermo Fisher Scientific, Kalamazoo, MI, USA).

### Droplet digital PCR

For RNA isolation from collected retinas, the Roche High Pure RNA kit (Roche Diagnostics GmbH, Mannheim, Germany; Cat. No. 11828665001) was used, and all procedures were carried out according to the manufacturer’s instructions. Complementary DNA (cDNA) was synthesized using the High-Capacity cDNA Reverse Transcription Kit (Applied Biosystems,Thermo Fisher Scientific, Waltham, MA, USA; Cat. No. 4368814) following the manufacturer’s instructions.

For droplet digital PCR (ddPCR), 0.25 ng of synthesized cDNA was used per reaction. The primers used in this study are listed in Table S1. All ddPCR steps including reaction setup, droplet generation, and amplification were conducted according to protocol as described in Okan et al ^48^.

### PBMC sample preparation

Blood samples were collected via cardiac puncture and diluted 1:2 with Hank’s Balanced Salt Solution (HBSS; Wisent Inc., Saint-Jean-Baptiste, QC, Canada; Cat. No. 311-512-CL). The diluted blood was carefully layered onto Ficoll-Paque™ PREMIUM density gradient medium (density, 1.084 g/mL; Cytiva Sweden AB, Uppsala, Sweden; Cat. No. 17-5446-02) at a Ficoll-to-diluted blood ratio of 3:4. Following density gradient centrifugation, the peripheral blood mononuclear cell (PBMC) layer was collected and subjected to the subsequent washing and centrifugation steps illustrated in Figure S1. The isolated PBMCs were then processed for downstream kinome profiling.

### Kinase activity analysis

Kinase activity analysis and upstream kinase analysis (UKA) were performed using PamChip® peptide microarrays on a PamStation®12 platform (PamGene International B.V., ‘s-Hertogenbosch, the Netherlands), as previously described ^49,50^, with minor modifications. Of the 51 PBMC lysates, 36 had detectable protein concentrations above 0.5 μg/μL. Serine/threonine kinase (STK) profiling was performed on all 36 samples using 1.5 μg of protein per array. Protein tyrosine kinase (PTK) profiling was performed on 34 samples using 5.0 μg per array, as insufficient lysate volume prevented PTK profiling of two LP-CN03-treated *rd10* samples. Four samples from each assay had no detectable kinase activity and were excluded from subsequent analyses. The final analyses included 11 wild-type, 11 untreated *rd10*, and 10 LP-CN03-treated *rd10* samples for STK, and 11, 11, and 8 samples, respectively, for PTK. Signal intensity data were subjected to variance stabilizing normalization (VSN) and ComBat correction for batch effects. Differences in peptide phosphorylation between groups were assessed using one-way ANOVA followed by Dunnett’s post-hoc test, with *p* < 0.05 considered significant. For UKA, a kinase score (median final score) > 1.3 was used as the default threshold for identifying relevant kinases.

The disease-related kinase profile was determined by comparing untreated *rd10* mice with wild-type controls (T1 *vs*. C). LP-CN03-treated *rd10* mice were compared with wild-type controls (T2 *vs*. C), whereas the treatment effect was examined by comparing LP-CN03-treated and untreated *rd10* mice (T2 *vs*. T1). Significant phosphosite and UKA results from each comparison were combined in a single input table and matched by UniProt accession. This table was used as input for pathway enrichment analysis, performed independently using the R package enrichR v3.4, using the 338 analysed kinases/substrates as background for enrichment analysis. Pathway and functional enrichment analyses were based on KEGG 2026 Human, Reactome Pathways 2024, GO Biological Process 2023, and WikiPathways 2024 Human. Terms were considered significant at a Benjamini–Hochberg adjusted *p-*value < 0.05.

## Statistical analysis

Statistical analyses were performed using Prism (GraphPad Software, Boston, MA), unless otherwise specified. The statistical tests, sample sizes, and presentation of data are indicated in the corresponding figure legends.

## Data Availability

Data supporting the findings of this study are available within the article and its supplemental information or from the corresponding author upon reasonable request.

## Acknowledgements

Generation of the *Aipl1*^-/-^ was carried out at the Izmir Biomedicine and Genome Center. Overall *in vivo* procedures were supported by the animal facilities at Acibadem Mehmet Ali Aydinlar University. Kinome profiling was performed by PamGene. Histological evaluations were conducted at the Virscio Pathology and Tissue Pharmacodynamics Laboratory. The patented LP-CN03 compound used in this study was provided via a collaboration with Mireca Medicines GmbH (Tübingen, Germany). The compound CN03 has received an orphan drug designation for retinitis pigmentosa (EU/3/15/1462), and its intellectual property is protected under patents WO2016146669A1 and US10322087B2 held by Mireca Medicines. Financial support for this work was provided by TUBITAK-BIDEB 2232 (118C226), the Integration Grant, Sabanci University, TUBITAK (121N275), the EJPRD TreatRP program, BMBF grant 16GW0267K, Sabanci University PRF (H-2021-0356), the Research Council of Kantonsspital Aarau (1410.000.188), Velux Stiftung (LMIC - 2012).

## Author Contributions

Conceptualization, C.A., F.P.-D. and M.K.D.; methodology, C.A., M.K.D., F.P.-D. and I.C.T.O.; investigation, I.C.T.O., M.A., H.Y.A., S.P.S., G.K., S.O., R.B., K.E., and M.K.D.; formal analysis, I.C.T.O., A.N., M.K.D, S.P.S., M.A., H.Y.A., G.K. and M.T.; resources, F.P.-D., P.G., M.T., and M.L.; visualization, I.C.T.O., M.K.D., A.N., H.Y.A. and S.P.S.; writing – original draft, I.C.T.O.; writing – review C editing, C.A., F.P.-D., M.K.D., A.N., M.T., M.A., H.Y.A. and S.P.S.; funding acquisition, C.A., F.P.-D., M.T., and A.N.

## Declaration of Interests

F.P.-D. (François Paquet-Durand) has filed for two patents related to the synthesis, use, and formulation of cGMP analogs (US10322087B2, US11407781B2). F.P.-D. and P.G. (Pieter Gaillard) hold shares in, or have other financial interests in, Mireca Medicines. S.P.S. (Simar Pal Singh) is an employee of PamGene International B.V. R.B. (Richard Bouffard), M.L. (Matthew Lawrence), and C.A. (Cavit Agca) are currently employed by Virscio. The remaining authors declare no competing financial interests.

