## Supplemental Information for "Identification of Kinome Biomarkers and Therapeutic Rescue in cGMP-Linked Retinal Degeneration"

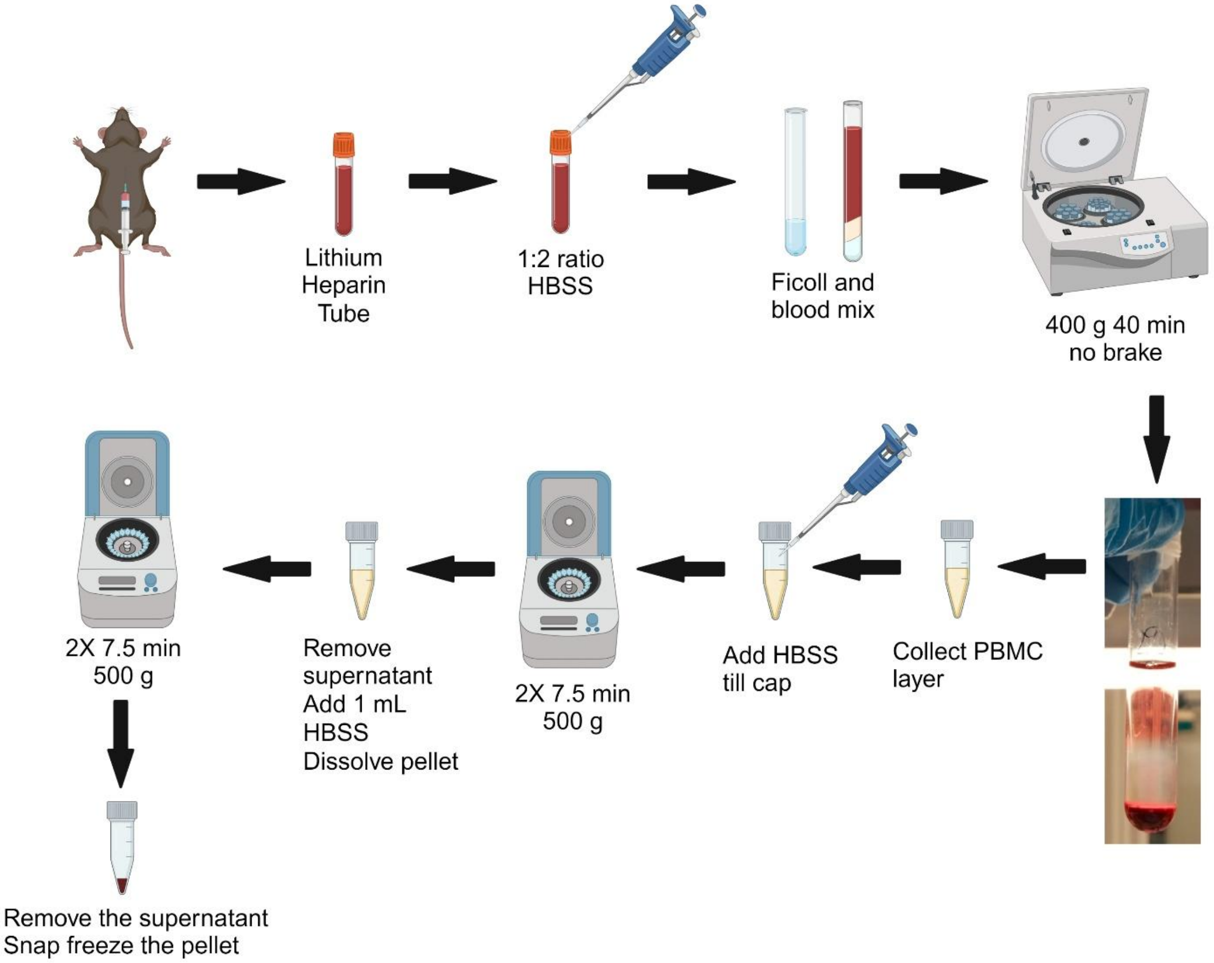


**Figure S1**. **Schematic representation of peripheral blood mononuclear cell (PBMC) isolation.** Blood samples were collected via cardiac puncture and diluted 1:2 with Hank's Balanced Salt Solution (HBSS). The diluted blood was carefully layered onto Ficoll density gradient medium at a Ficoll-to-diluted blood ratio of 3:4. Following density gradient centrifugation, the PBMC layer was collected and subjected to two washing and centrifugation steps. The isolated PBMCs were then processed for downstream kinome profiling. Created in BioRender. Okan, I. C. T. (2026) https://BioRender.com/58hbxod.

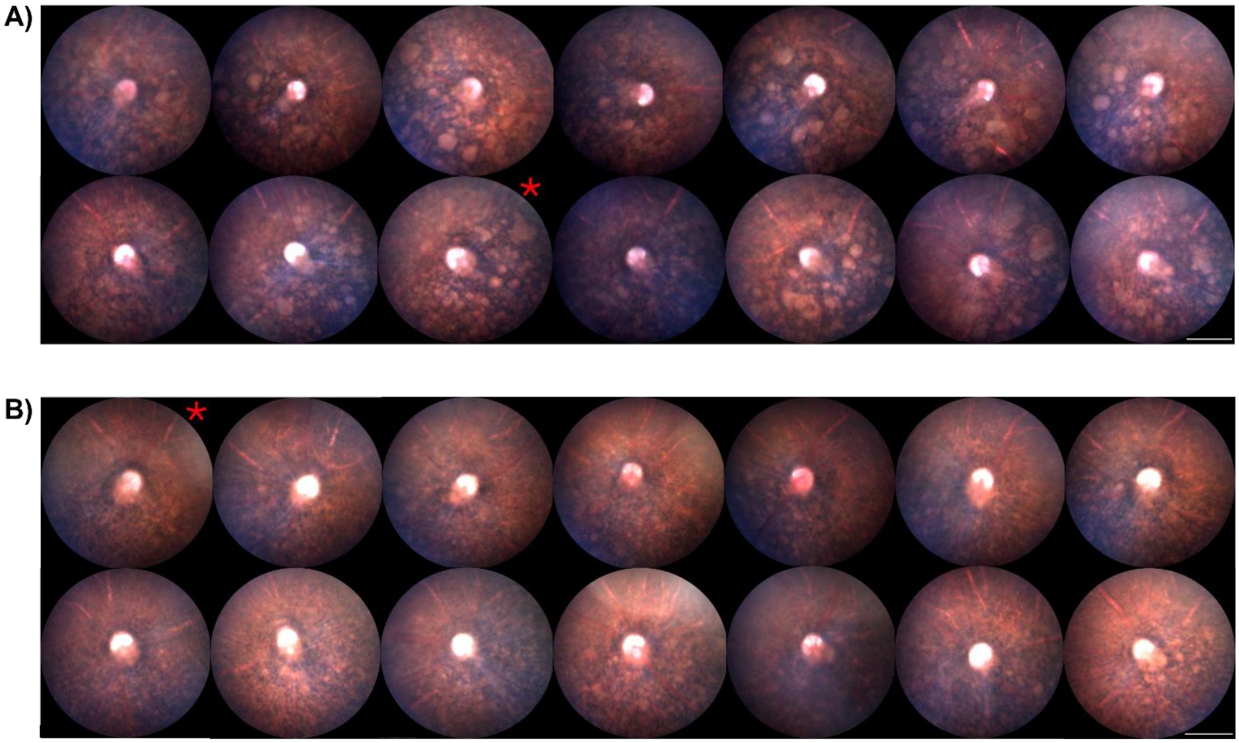


**Figure S2. Effect of LP-CN03 treatment on fundus appearance in *Aipl1*$^{-/-}$ mice.** Fundus images of untreated **A**) and LP-CN03-treated *Aipl1*$^{-/-}$ mice **B**) at post-natal day 23. An apparent reduction in multifocal fundus abnormalities and pigmentary mottling is observed in treated animals compared with untreated controls. Red asterisks indicate the animals shown in Figure 6A. Scale bars represent 500 µm.

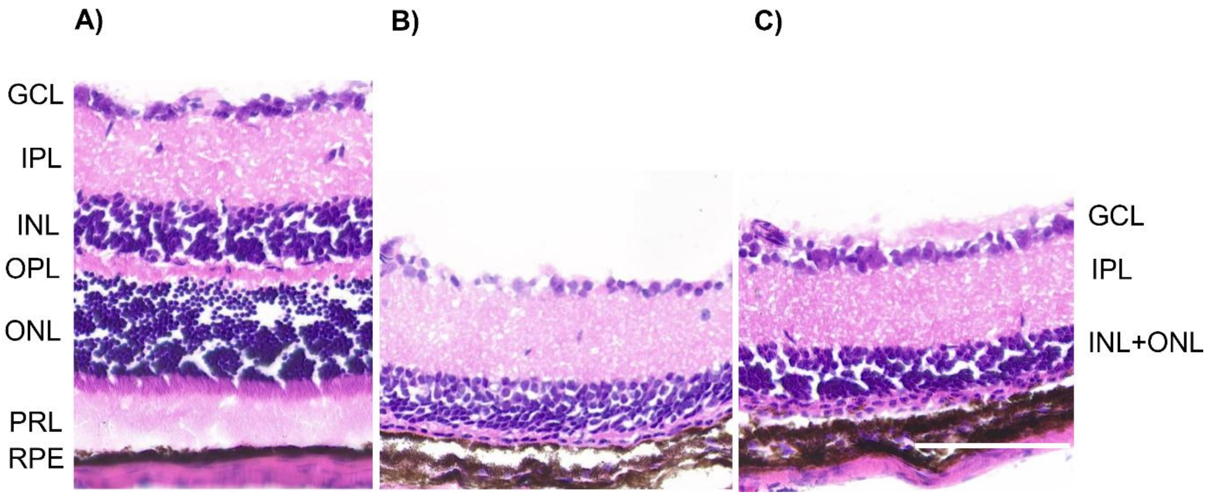

**Figure S3. Progression of photoreceptor loss in *Aipl1*$^{-/-}$ mice.** Representative haematoxylin & eosin staining of retinal sections derived from P 24 **A**) wild-type, **B**) untreated *Aipl1*$^{-/-}$, and **C**) LP-CN03 treated *Aipl1*$^{-/-}$ mice. Scale bar represents 100 µm.

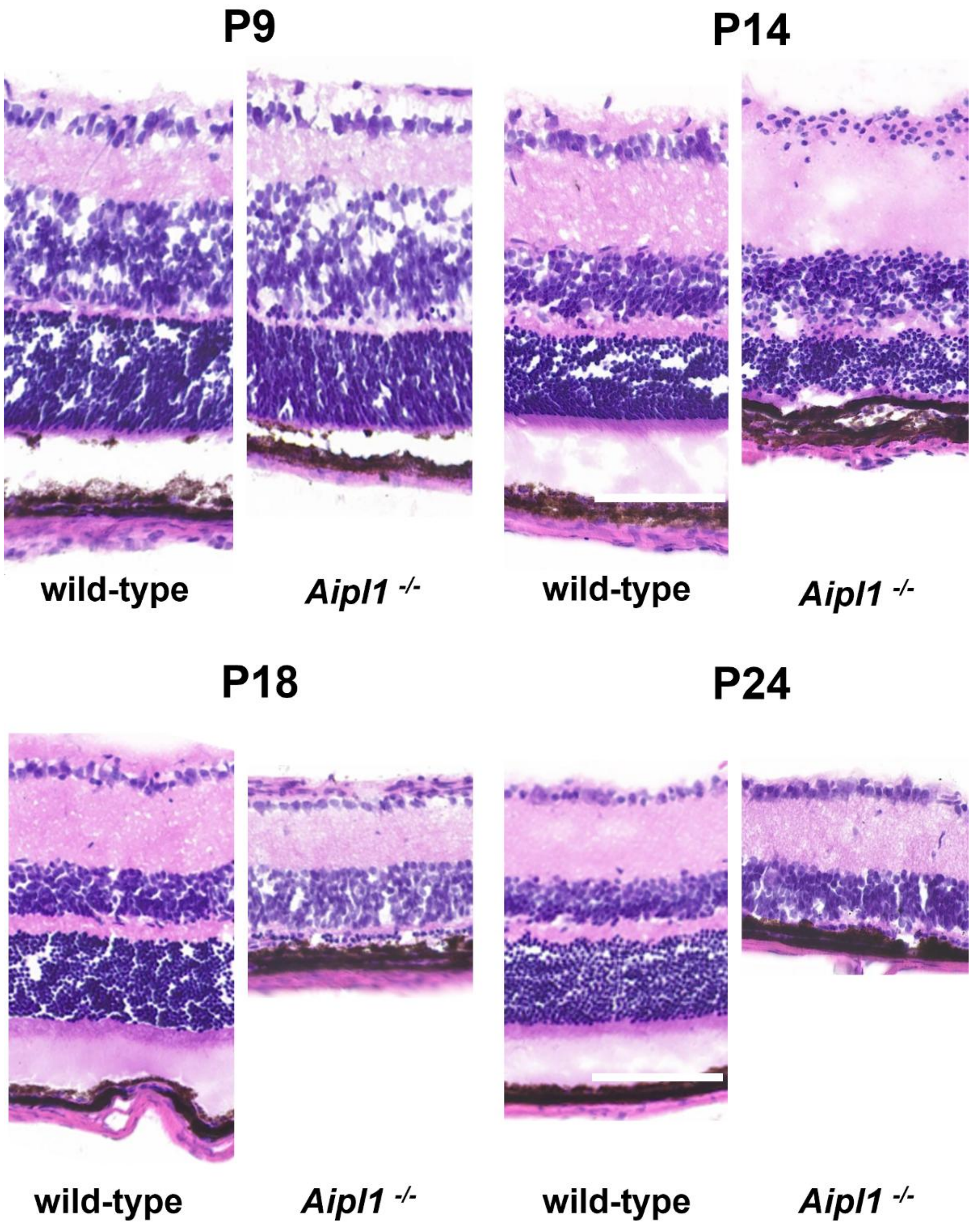


**Figure S4. Comparative analysis of disease progression in *Aipl1*$^{-/-}$ mice.** Representative haematoxylin and eosin stained retinal sections from age-matched wild-type (left) and *Aipl1*$^{-/-}$ (right) mice at post-natal days 9,

14, 18, and 24. Images illustrate the progressive loss of the photoreceptor layer in *Aipl1*$^{-/-}$ retinas. Scale bars represent 100 µm.

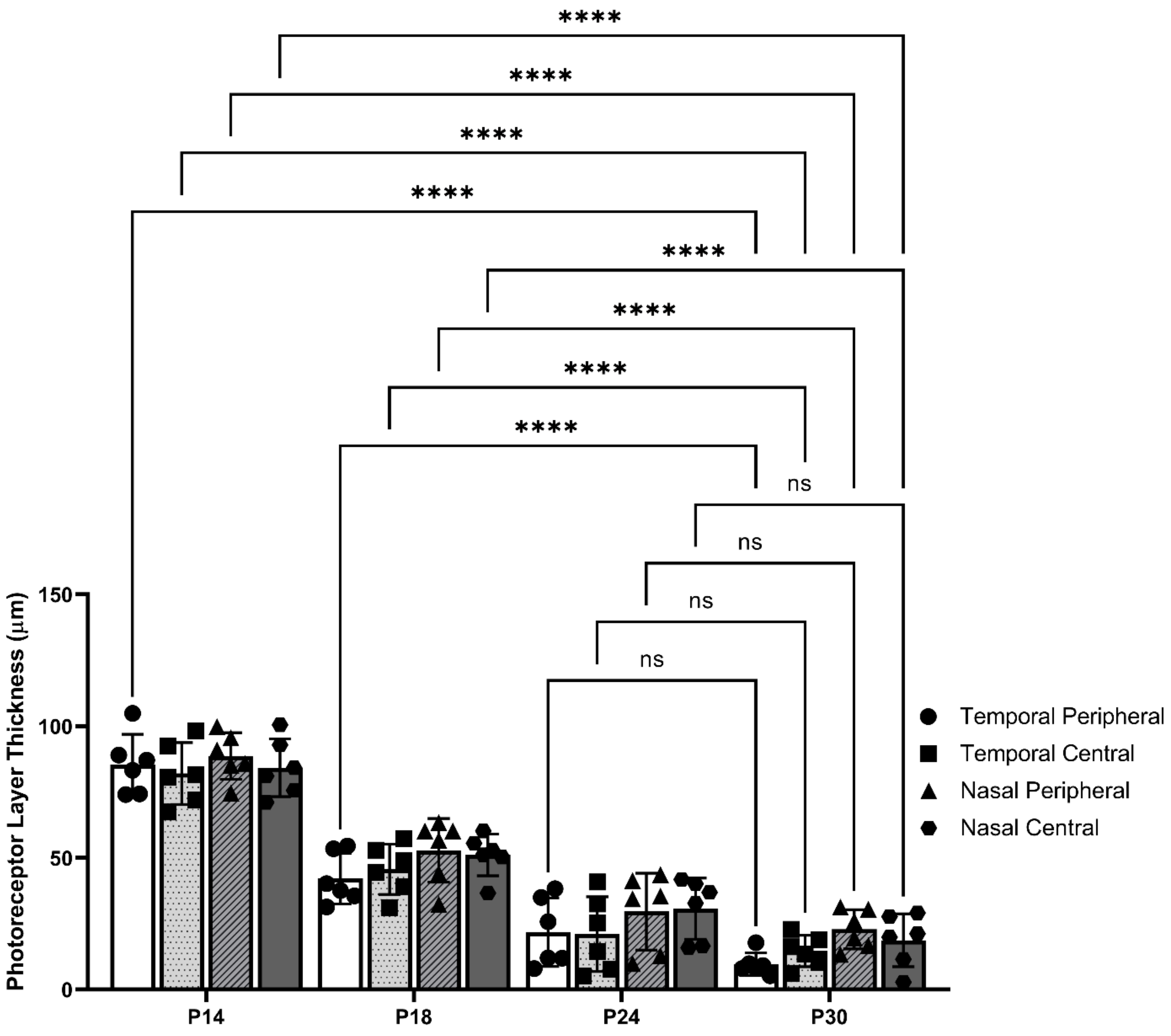


**Figure S5. OCT analysis of photoreceptor layer thickness during retinal degeneration in *Aipl1*$^{-/-}$ mice.** Photoreceptor layer thickness was measured from OCT images acquired at post-natal day 14, 18, 24, and 30 at four retinal locations corresponding to the temporal peripheral, temporal central, nasal peripheral, and nasal central regions. Data are presented as mean ± SD (n = 6 eyes from 3 animals per time point). Statistical significance was determined using ordinary two-way ANOVA followed by Tukey's multiple comparisons test. ns, not significant; $^{**}p < 0.01$; $^{***}p < 0.001$; $^{****}p < 0.0001$.

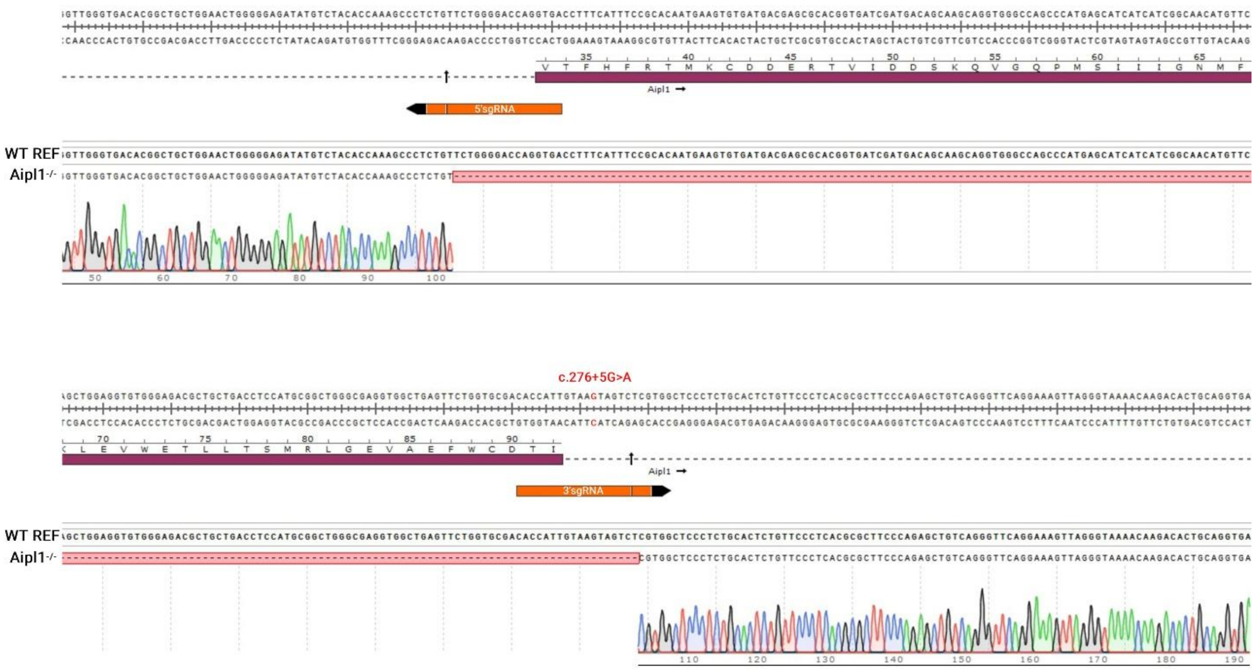


**Figure S6. Sanger sequencing analysis of the *Aipl1* exon 2 deletion allele.** PCR products from two independent homozygous *Aipl1*$^{-/-}$ mice were analyzed by Sanger sequencing, with a representative chromatogram from one of these mice presented here. The resulting sequencing chromatogram is aligned to the WT reference sequence, illustrating a 203 bp deletion including exon 2 generated by CRISPR/Cas9-mediated genome editing. Up-pointing arrows indicate the locations of the CRISPR/Cas9-generated DNA double-strand breaks (DSBs). Additionally, the locations of exon 2, the sgRNA target sites, and the *Aipl1*$^{tvrm119}$ mutation (c.276+5G>A) are indicated.

**Table S1. Droplet digital PCR Primer List.**

| Primer List for ddPCR | |
|---|---|
| *Gapdh* Forward | CAGCAATGCATCCTGCACC |
| *Gapdh* Reverse | TGGACTGTGGTCATGAGCCC |
| *Rho* Forward | CTTCACCTGGATCATGGCGTT |
| *Rho* Reverse | TCGTTGTTGACCTCAGGCTT |
| *Opn1mw* Forward | CTCTGCTACCTCCAAGTGTGG |
| *Opn1mw* Reverse | AAGTATAGGGTCCCCAGCAGA |
| *Opn1sw* Forward | TGTACATGGTCAACAATCGGA |
| *Opn1sw* Reverse | ACACCATCTCCAGAATGCAAG |
| *Gnat1* Forward | GAGGATGCTGAGAAGGATGC |
| *Gnat1* Reverse | TGAATGTTGAGCGTGGTCAT |
| *Gnat2* Forward | GCATCAGTGCTGAGGACAAA |
| *Gnat2* Reverse | CTAGGCACTCTTCGGGTGAG |

**Table S2. Genotyping and Sequencing Primers**

| PCR Primers | |
|---|---|
| FOR1 *(Aipl1)* | AGGCATAGGGTTTGCTGGAA |
| FOR2 *(Aipl1)* | AGGGGAGCTGAGGCATCTTA |
| REV *(Aipl1)* | CACGTCACTTTAAGGCCTGC |
| *rd10* Forward *(Pde6b)* | CTTTCTATTCTCTGTCAGCAAAGC |
| *rd10* Reverse *(Pde6b)* | CATGAGTAGGGTAAACATGGTCTG |

**Table S3. Scotopic Electroretinography Protocol.**

| | - Intensity series, Scotopic 0 using a 1:1000 (Neutral Density 3) filter - | | | | | |
|---|---|---|---|---|---|---|
| **Step label** | **Action** | **Flash Intensity Log** | **mcd.s/m$^2$** | **Number of Flashes** | **Interval Seconds** | **Time Required Seconds** |
| T1_7 | Test 1 | -5 | 0.03 | 10 | 2 | 18 |
| T2_7D | Delay | | | | | 60 |
| T2_7 | Test 2 | -4.5 | 0.1 | 10 | 2 | 18 |
| T3_7D | Delay | | | | | 60 |
| T3_7 | Test 3 | -4 | 0.3 | 10 | 2 | 18 |
| T4_7D | Delay | | | | | 60 |
| T4_7 | Test 4 | -3.5 | 1.0 | 4 | 2 | 6 |
| T5_7D | Delay | | | | | 60 |
| T5_7 | Test 5 | -3 | 3.0 | 4 | 2 | 6 |
| T6_7D | Delay | | | | | 60 |
| T6_7 | Test 6 | -2.5 | 10 | 4 | 2 | 6 |
| T7_7D | Delay | | | | | 60 |
| T7_7 | Test 7 | -2 | 30 | 4 | 10 | 30 |

**Table S4. Phosphosite and upstream kinase analysis results from PTK and STK assays.** Provided as a separate Excel file.
